# Expanding the diversity of *Accumulibacter* with a novel type and deciphering the transcriptional and morphological features among co-occurring strains

**DOI:** 10.1101/2022.12.09.519852

**Authors:** Zhongjie Wang, Wei Song, Xue Zhang, Minjia Zheng, Hao Li, Ke Yu, Feng Guo

## Abstract

*Accumulibacter* is the major polyphosphate-accumulating organism (PAO) in global wastewater treatment systems. Its phylogenetic and functional diversity has been continuously updated in recent years. In addition to its widely recognized two sublineages, Types I and II, here we discovered a novel type enriched in laboratory bioreactors. Core gene- and machine learning-based gene feature profiling supported that Type III *Accumulibacter* was potential PAO with the unique function of using dimethyl sulfoxide as electron acceptor. On the basis of the correlation between the similarity of *ppk1* and genome, the number of *ppk1*-represented *Accumulibacter* species was estimated to be over one hundred, suggesting that the currently recognized species are only the tip of the iceberg. Meanwhile, multiple *Accumulibacter* strains co-occurring in a bioreactor were investigated for their inter-strain transcriptional and morphological features. Metatranscriptomics of seven co-occurring strains indicated that the expression level and interphasic dynamics of PAO phenotype-related genes had minimal correlation with their phylogeny. In particular, expression of denitrifying and poly-P metabolism genes had higher inter-strain and interphasic divergence compared with glycogen and polyhydroxyalkanoates metabolic genes. A strategy of cloning rRNA genes from different strains based on similar genomic synteny was successfully applied to differentiate their morphology via fluorescence in situ hybridization. Our study further expanded the phylogenetic and functional diversity of *Accumulibacter*. We also proposed that deciphering the function and capability of certain *Accumulibacter* should be environment- and population-specific.

**Importance:** *Accumulibacter*, as the core functional but uncultured taxa for enhanced biological phosphorus removal, has attracted much attentions on its phylogenetic and functional diversity and intra-genus niche differentiation in the last two decades. It was well-known that this genus had two sub-lineages (Type I and II) since 2002. In this study, a novel type (Type III) with proposed novel functional feature was discovered by the metagenomic approach. By linking average nucleotide identity of *Accumulibacter* genomes and the similarity of the *ppk1* sequences, a phylogenetic biomarker that has been largely deposited in database, we estimated that its global species-level diversity was higher than 100. Moreover, as we found the co-occurrence of multiple *Accumulibacter* strains in one bioreactor, the simultaneous transcriptional divergence of the co-occurring strains was interesting for understanding their niche differentiation in a single community. The results suggested the decoupling feature between transcriptional pattern with phylogeny for co-occurring strains.

## Introduction

Enhanced biological phosphate removal (EBPR) for wastewater treatment essentially relies on polyphosphate-accumulating organisms (PAOs) that can accumulate excessive phosphate within their cells (1–3). Typical PAOs have a metabolic shift that uptakes volatile fatty acid and generates energy from polyphosphate (poly-P) de-polymerization and glycogen glycolysis for polyhydroxyalkanoate (PHA) biosynthesis during the anaerobic phase. Then, glycogen and polyphosphate are replenished by using the energy produced from PHA degradation and conducting TCA cycling during the aerobic phase. The most prevalent and well-studied PAO in laboratory enrichments and full-scale activated sludge is *Candidatus* Accumulibacter (*Accumulibacter* hereafter), an uncultured genus affiliated with *Rhodocyclaceae*. On the basis of the polyphosphate kinase 1 gene (*ppk1*), *Accumulibacter* has been phylogenetically divided into two sublineages, namely, Type I and II, each of which contains multiple clades. To date, diverse clades have been proposed, i.e., Clade IA to IF and Clade IIA to II–I (4–6).

In the last two decades, *Accumulibacter* has been extensively studied by meta-omic approaches (7–10). Clade IIA strain UW-1 is the most well-studied organism that greatly contributes to understanding the metabolic paradigm of PAOs. On one hand, different carbon sources, electron acceptors, DO levels, phosphate concentration, and inoculum (11–15), have strong selectivity on the enrichment of certain types or clades. On the other hand, comparative genomics revealed their core and characteristic functions in carbon, nitrogen, and phosphorus metabolisms (16, 17). Both aspects indicated a specialized ecological adaption or niche differentiation among species or even strains. In addition, various *Accumulibacter* types, clades, species, and strains often co-occur in full-scale plants or laboratory-scale bioreactors (14, 18, 19). McDaniel *et al*. (18) recently reported the simultaneously transcriptomic divergence of co-occurring *Accumulibacter* strains (focusing on two strains although four were detected) and deduced the potential niche differentiation of corresponding strains.

Understanding the phylogenetic and functional diversity of *Accumulibacter* is significant for promoting EBPR performance. In this study, we aim to further explore the genomic diversity of *Accumulibacter* and investigate the transcriptional and morphological features of co-occurring *Accumulibacter* strains. Laboratory bioreactors were operated by creating diverse niches, such as using acetate and propionate as dual carbon sources and allowing full nitrification/denitrification, to set up *Accumulibacter* enrichments containing multiple clades. Genomes affiliated with *Accumulibacter*, including those representing a novel type, were then retrieved for genomic analysis. On the basis of the good phylogenetic correlation between genomes and *ppk1*, the global species-level diversity of *Accumulibacter* was estimated. For co-occurring *Accumulibacter* strains, the expression dynamics of key genes related to EPBR performance was compared based on genome-resolved metatranscriptomics. Finally, fluorescence *in situ* hybridization (FISH) based on a specific strategy was conducted to obtain the full-length 16S rRNA gene sequences of co-occurring *Accumulibacter* strains for strain-level morphological characterization.

## Materials and Methods

### Operation of EBPR bioreactors and sample collection

Two 3-L EBPR bioreactors (R1 and R2) were operated in the laboratory from May 19, 2016 to October 15, 2018. The basic setting in each cycle was 2 h under anaerobic conditions (with stirring at 50 rpm) after the introduction of influent, 3 h under aerobic conditions (aeration rate at 3 L/min), 50 min of settling, 5 min of discharge, and 5 min of pumping influent. The influent contained acetate (300 mg L^-1^ COD) and propionate (300 mg L^-1^ COD) as carbon sources. Nitrification and denitrification were facilitated by adding 1 mM NH_3_-N (NH_4_Cl) to the influent. Other operational information can be found in our previous report (14). The only difference between the two bioreactors was the aeration strength: per minute 3- and 0.5-liter air was pumped in to R1 and R2, respectively, during the aerobic phase. The dissolved oxygen (DO) of R1 and R2 during the aerobic phase typically ranged at 4–8 and 0.5–2 mg L^-1^, respectively.

DNA samples for R1 and R2 were collected on March 23, 2017. In the exact cycle, biomass samples from R1 were collected for metatranscriptomic analysis at the 15th min (anaerobic phase, designated as AN) and 130th min (aerobic phase, designated as AE). FISH samples were collected on October 15, 2018. Water samples collected at 5, 15, 35, 55, 85, 110, 130, 145, 175, 205, 235, 265, 325, and 350 min of the exact operational cycle were examined for DO, pH, NH_3_-N, NO_3_-N, COD, and PO_4_-P. Replications of biomass samples were fixed in 4% paraformaldehyde (for FISH, stored at 4 °C), 50% ethanol (for DNA extraction, stored at −20 °C), and RNALater (for RNA extraction, stored at −80 °C).

### DNA and RNA extraction

DNA and RNA were extracted by using FastDNA SPIN Kit for Soil (MP Biomedicals, US) and PowerSoil Total RNA Isolation Kit (QIAGEN, Germany), respectively. Their quantity and quality were determined with a microspectrophotometer (OD260/280 >1.7 for DNA and 1.9 for RNA). RNA integrity was further examined by Bioanalyzer device (RIN value >6.5, Agilent 2100, USA).

### High-throughput sequencing

Metagenomic and metatranscriptomic sequencing were conducted in Hiseq X10 platform using the PE150 strategy. For metatranscriptomic sequencing, total RNA was cleaned by genomic removal and then reversely transcribed (Prime Script RT reagent Kit with gDNA Eraser, Takara, China). The rRNA removal, library construction, and sequencing services were provided by Novogene (Beijing, China). Clean data were delivered after standardized QC criterion. Approximately 19.6 G and 20.2 G data were obtained for the R1 and R2 metagenomes, respectively. 41.7 G and 53.6 G data were obtained for AN and AE metatranscriptomes, respectively.

### Genomic binning and annotation

Genome binning was performed using the toolkit BASALT (20), which assembles multiple sample files in parallel before incorporating custom algorithms to separate redundant bins for refinement. It can further calibrate assembled bins and complement gaps using third-generation sequencing data. After genome qualities were estimated using CheckM v1.1.3 (21), *Accumulibacter* genomes (requiring completeness >70%, contamination <10%) were retained for further analysis. Four genomes were used as outgroup (*Rhodocylcus tenuis* DSM109, *Dechloromonas hortensis* DSM15637, and two *Propionivibrio* genomes). Besides, 67 *Accumulibacter* genomes publicly available online were downloaded from the NCBI genome database (Aug 20th, 2021). For all the above genomes, open reading frames (ORFs) were predicted using Prodigal v2.6.3 (22). Genome metabolic functions were annotated using GhostKOALA (23).

### Phylogenetic analysis and gene feature mining

The *ppk1* extracted from *Accumulibacter* genome bins and the reference sequences (5, 24) were used to infer the phylogenetic tree in MEGA X (25). The set of 120 bacterial conserved single-copy marker genes (bac120) was extracted from the genomes, aligned, and trimmed following the method of Parks et al.(26). A phylogenetic tree was then constructed on the basis of the concatenated bac120 genes with a model of PROTGAMMAAUTO in RAxML v8.2.11 with 200 bootstraps (27). All trees were visualized and further polished using the interactive tree of life (iTOL) v5 (28).

All-against-all BLASTP for all genomes was carried out with the parameter -evalue 1e-5 to determine the core gene catalog for *Accumulibacter* (29). The result was then filtered to percent identity of 70% and query coverage of 75% (30). Finally, orthologous gene clusters were identified using MCL with an inflation value of 2 (31). Core gene set was determined by referring to the method of Oyserman et al. (2016a) for *Accumulibacter* Type I and II. However, due to the great number of genomes costing excessive time and computational resources, the calculation was terminated at an appropriate time point when approximately 99% of core genes could be identified by applying a cutoff. Core genes were defined as those present in the number of genomes above the cutoff (n=58 among the 68 genomes).

Random Forest (RF) machine learning algorithm based on KEGG annotation results was adopted to determine the key gene features of *Accumulibacter* and outgroup. The dataset was randomly split into a training dataset (59 out of 65 Types I and II *Accumulibacter* genomes, five out of six outgroup genomes) to build the classifier model and a validation dataset (six genomes from Type I or II *Accumulibacter* and one genome from outgroup), which was repeated 50 times. The RF classification model was first trained using the R package RandomForest v4.6-14 (32) with the parameters of 501 trees (ntree=501) and the default number of features randomly sampled at each node in a tree (mtry). Finally, medians of mean decrease in accuracy of the 50 repetitions were calculated to determine the top 20 important gene features.

### Estimation of species-level diversity of *Accumulibacter* via linking genomic average nucleotide identity and sequence similarity of *ppk1*

Seed sequences (ref(5)) were collected to extract all *ppk1* from *Accumulibacter* from the NCBI nucleotide (NT) database. The top 1,000 hits for each seed sequence were downloaded for further curation. They were first cleaned by de-replication and length (>1,000 bp). The remaining unique *ppk1* genes were aligned using MAFFT v2.6.3 (33), trimmed using trimAl v1.4.43v22 (34), and further processed for *de novo* operational taxonomic unit (OTU) clustering in Mothur v1.39.5 (35) with the furthest, nearest, and average algorithms under various similarity thresholds (100%–91%, 1% stepwise decrease). The representative sequences of OTUs with the furthest algorithm and cutoff of 91% were selected to construct a phylogenetic tree, and the *ppk1* sequences not derived from *Accumulibacter* were removed. The *ppk1* sequences represented by the remaining OTUs were used to estimate the species level diversity of *Accumulibacter* by repeating the OTU clustering. For all *Accumulibacter* genomes with *ppk1*, the pairwise distance of *ppk1* and the genomic average nucleotide identity (ANI) were calculated in MEGA X (25) and FastANI v1.33 (36), respectively. A species-level threshold was then determined for the *ppk1* similarity, and the global species-level diversity of *Accumulibacter* was calculated using the all cleaned *Accumulibacter ppk1* sequences.

### Genome-centered metatranscriptomic analysis

The expression data of *Accumulibacter* was calculated by mapping the metatranscriptomic data to the genome bins using Bowtie2 (37) and Samtools (38). Since relative expression level of a given gene for a strain was concerned, gene expression was calculated as: reads per million (RPM) of a gene = (number of reads mapped to a gene in a given genome × 10^6^) / (total number of mapped reads from a given genome). The genes involved in denitrification, PHA metabolism, glycogen metabolism, and polyphosphate (poly-P) accumulation were singled out to compare their expression variations among the co-occurring *Accumulibacter* strains under anaerobic and aerobic phases. The results were then visualized using pheatmap v 1.0.12 (39).

### Retrieving the full-length 16S rRNA gene for two *Accumulibacter* strains and their FISH detection

Due to the absence of rRNA gene in all *Accumulibacter* bins, a strategy was adopted to clone the 16S rRNA gene sequence for the two most abundant bins. We hypothesized that the genomic location of rRNA operon is conserved for the two bins and UW-1, a Clade IIA strain with a complete genome. Therefore, the homogeneous region of the two bins corresponding to the upstream region of one 16S rRNA gene copy in the UW-1 genome was first identified. Fortunately, the homogeneous region, which showed breaking near the 16S rRNA gene, was found in both bins. A forward primer was then designed at the end of the upstream homogeneous region for each bin. PCR was performed by using the forward primer and 1492R, the universal primer for bacterial 16S rRNA gene. The 16S rRNA genes of the two bins were successfully cloned and sequenced. Finally, unique probes were designed on the basis of the two sequences (mostly divergent region was selected to discriminate the two bins). In addition to the bin-specific probes, EUB338-mix and PAO-mix probes respectively targeting most bacteria and all *Accumulibacter* were applied. FISH was performed as previously described (40), and the FISH samples were examined with a confocal microscope.

## Results

### Operational information for the EBPR bioreactors

A typical nitrifying–denitrifying EBPR performance was observed for the COD and PO_4_-P of the reactors (see data for the cycle of metatranscriptomic sample collection in Fig S1). COD rapidly decreased to approximately 80 mg/L during the first few minutes after the injection of influent and then remained at 40–60 mg/L. Phosphate in the medium increased from 23 mg/L to around 40 mg/L in the early anaerobic phase and dramatically decreased to an almost undetectable level within 1 h after initial aeration. NH_4_-N was at 4–5 mg/L during the initial anaerobic phase and rapidly depleted during the aerobic phase, which should be mainly attributed to ammonia oxidization. Interestingly, NO_3_-N was maintained at about 2 mg/L during the aerobic phase, which was in imbalance with the stoichiometry of NH_3_-N decrease because of the lack of strong NO_2_-N accumulation (<0.2 mg/L in all samples, data not shown). This phenomenon might have resulted from the denitrification by anaerobic denitrifiers whose microenvironment was anaerobic during the aerobic phase or by some aerobic denitrifiers. The occurrence of denitrification was supported by the pH increasing shortly after aeration (from 7.2 at 110 min to 8.5 at 130 min). For NO_3_-N, its high concentration in the last sample (350th min) and low concentration in the first sample (5th min) should be attributed to expeditious denitrification during the injection of fresh medium (lasting 5 min, supported by the fact that COD decreasing rapidly), although the pH also decreased possibly by the buffering effect from the high phosphate concentration. The two metatranscriptomic samples (AN and AE) were collected at 15 and 130 min. AN showed a sharp COD decrease and P-efflux, and AE exhibited rapid P-influx and the emergence of NO_3_-N.

### Retrieving 10 *Accumulibacter* genomes including one novel clade (Clade IG) and one novel type (Type III) from metagenomes

A total of 102 dereplicated and quality-filtered bins (completeness-5×contamination>50%) were retrieved from the two metagenomic datasets by genome binning for R1 and R2 metagenomes. Their taxonomic information was determined by GTDB-Tk (see detailed information in Table S1). Genomes that can be classified as *Accumulibacter* and unclassified *Rhodocyclaceae* were selected. For the latter ones, their *ppk1* sequences were extracted and searched against Genbank. Once the best hit was an *Accumulibacter ppk1*, the corresponding genome was also thought to be potentially affiliated with *Accumulibacter*. Ten candidate *Accumulibacter* bins (i.e., Bin28, 45, 87, 98, 136, 140, 142, 202, 208, and 228) were retrieved from the metagenomes with completeness and contamination ranging 78.5%–99.5% and 0–3.8%, respectively (Table S1). Only two genomes, Bin87 and Bin228, belong to a single species (ANI=97%, Fig S2). The genome coverages for the 10 *Accumulibacter* strains in the two metagenomes of R1 and R2 are shown in Fig S3. Two glycogen-accumulating organism (GAO) genomes affiliated with *Competibacteraceae* were also retrieved with high quality (Table S1).

The phylogeny was reconstructed on the basis of the *ppk1* sequences to assign the 10 *Accumulibacter* genomes into types and clades. We successfully extracted this single-copy gene from all bins except Bin142. A *ppk1* phylogenetic tree was reconstructed by combining the genome-derived *ppk1* sequences with representative references (5, 24). Figure 1A shows that in addition to the known 15 clades (Clade IA to IF and IIA to II–I), two novel lineages were identified and proposed as Clade IG (Bin208 and three reference sequences) and Type III (Bin87 and Bin 228) considering their remote phylogenetic distance to the previously defined clades and types. Bin28, 45, 98, 136, and 202 were assigned to Clades IIC, IIA, IIB, IIF, and IID, respectively. Bin140 represented the recently proposed Clade IF (6). Noticeably, primer mismatch was observed for the two Type III genome-derived *ppk1* sequences. Mismatches occurred near the 3’ end of the primer 254F (Fig S5), indicating that PCR is unlikely successful by using the primer.

**Figure 1.**
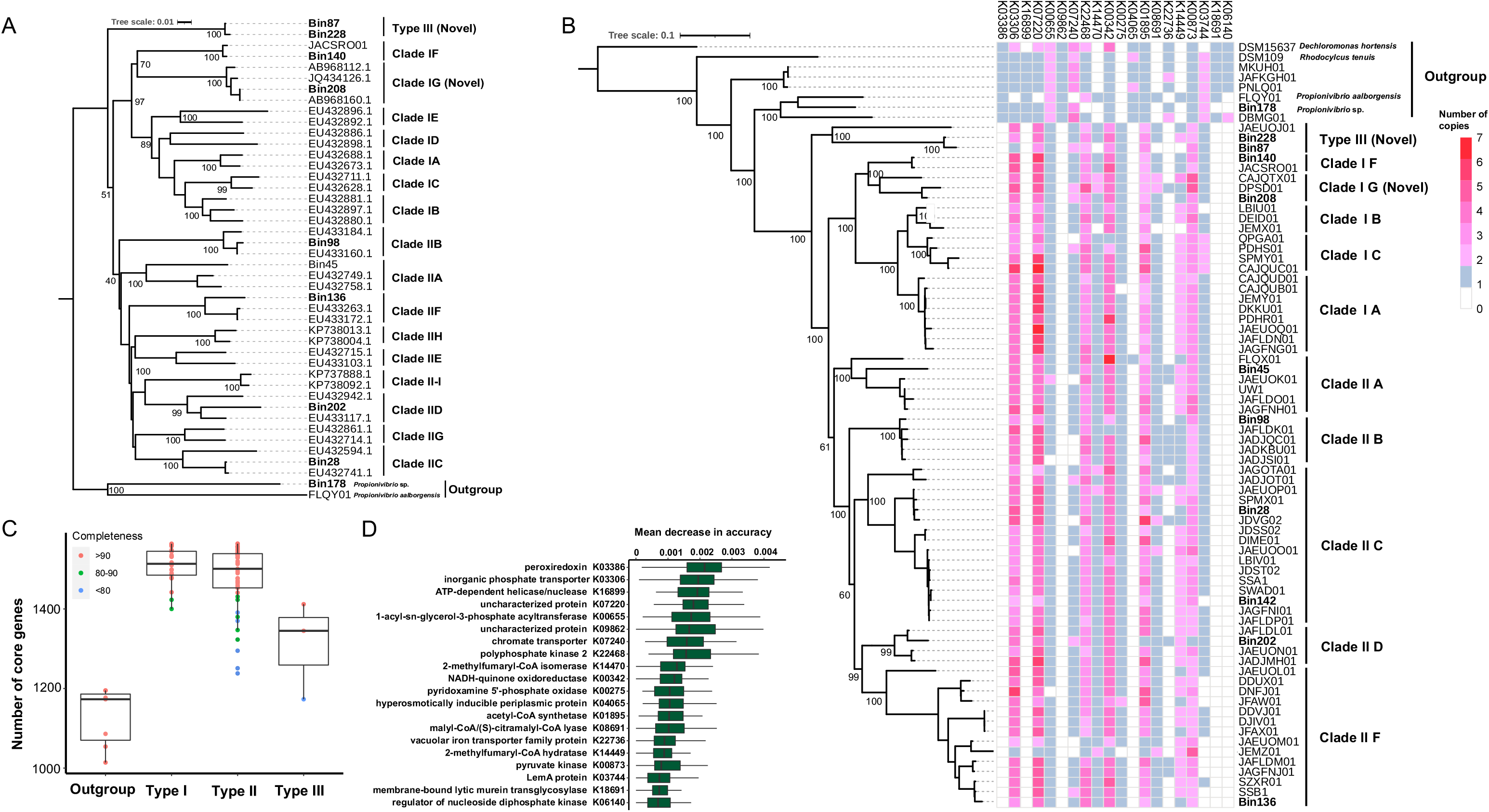
Phylogenetics and gene feature of *Accumulibacter*. Phylogenetic trees were reconstructed based on both *ppk1* (A) and the concatenated bac120 genes (B). Bootstrapping (500 times for neighbor-join tree of *ppk1* and 200 times for maximum-likelihood tree of bac120) was performed to find the best topology. Bootstrap values were shown beside the nodes of types and clades. Genome bins extracted from metagenomes in this study were marked in bold. Outgroups were introduced in both phylogenetic trees. Reroot was conducted only for the bac120 tree. The number of core genes of outgroup and types of *Accumulibacter* was shown in C. The mean decrease accuracy value for the top 20 gene features differentiating *Accumulibacter* from outgroups identified with the random forest model were shown in D.

The phylogenomics of 68 high-quality *Accumulibacter* genomes (information on the reference genomes can be found in Table S2) obtained by concatenating the alignments of bac120 genes is shown in Figure 1B. Overall type-level and clade-level consistency between phylogenomics and *ppk1*-based phylogeny was confirmed. The two proposed novel lineages (Clade IG and Type III) were supported by their phylogenomic distances from other defined clades and the strength of the corresponding nodes (Figure 1B). Three reference genomes were also assigned to the two novel lineages (CAJQTX and JACSRO in Clade IG and JAEUOJ in Type III). This finding suggested that the novel lineages may not be occasionally present in our EBPR systems. For Bin142, which has no detected *ppk1*, phylogenomics assigned it as a IIC strain. Therefore, the 10 *Accumulibacter* genomes represented eight clades including a novel clade and a novel type.

### Comparative genomics supports that Type III *Accumulibacter* strains are PAOs with unique electron acceptor

Given that the Type III strains are phylogenetically located outside Types I and II (Figures 1A&B), their functions predicted by genome features, especially those related to PAO phenotypes, require further investigation. First, we profiled the distribution of core genes defined by Type I and II genomes in the Type III genomes and outgroups. Type III genomes had slightly fewer core genes than Types I and II but much higher ones than outgroup members (Fig 1C). The missing functions (defined as core genes representing unique KEGG functions that are not detected in any Type III genomes) were only eight, and none of which are putatively involved in poly-P accumulating phenotype (Table S3). We also performed the machine learning-based profiling of key feature genes to differentiate known *Accumulibacter* (Type I and II) from outgroups. For the top 20 features (Fig 1D), Type III genomes showed a similar distribution pattern to Type I and II genomes (heatmap in Figure 1B). Three key PAO-related metabolic genes (K03306, *Pit*, low-affinity inorganic phosphate transporter; K22486, *ppk2*, poly-P kinase2; K01895, *acs*, acetyl-CoA synthetase) were detected in all *Accumulibacter* and outgroups, and *Accumulibacter* (including Type III) have multiple copies in contrast to the single-copy feature in the nearly all of the outgroup members.

We then profiled the unique functional genes present in the Type III genomes (detected in at least two of the three Type III genomes) but absent in all Type I and II genomes. Only five of these Type III specific genes, i.e., K07306, K07307, K07131, K02849, and K06075, were identified. K07306 and K07307, which were found in all three Type III genomes, are anaerobic dimethyl sulfoxide (DMSO) reductase subunit A and B, respectively. K07131 is functionally unknown. K02849 and K06075 are heptosyltransferase III and MarR family transcriptional regulator, respectively. The niche adaptation endowed by the latter three genes is not clear. However, the presence of anaerobic DMSO reductase indicates a novel organic electron acceptor that has not been proposed in *Accumulibacter*. In addition, the denitrification pathway is incomplete in all Type III genomes and other *Accumulibacter* strains except for the Clade IC strain QPGA01 (information on the distribution of key genes related to phosphate accumulation, denitrification, and PHA and glycogen metabolism can be found in Table S4).

### Estimation of global species-level diversity of *Accumulibacter*

The global species-level diversity of *Accumulibacter* has not been estimated due to the lack of sufficient genomes, despite the large numbers of *ppk1* sequences deposited in GenBank. Studies, including the present one (Figures 1A & B), suggested that the phylogeny of *ppk1* has a robust consistency with phylogenomics for *Accumulibacter*. With the accumulation of *Accumulibacter* genomes, a species-level *ppk1* threshold for *Accumulibacter* may be proposed by correlating the distances of *ppk1* and ANI values between genome pairs. A good negative correlation between the distance of *ppk1* and ANI values was observed (R^2^=0.9545, Figure 2A). On the basis of this correlation, the species-level threshold of *ppk1* distance was set at 0.04. Hence, the predicted species number for the 63 genomes with *ppk1* was 22 (the furthest clustering algorithm), which is slightly lower than 26, the true species-level diversity determined by ANI. We found that the few outliners corresponding to the threshold were contributed by four non-conspecific strains in Clade IC (QPGA01, CAJQUC01, PDHS01, and SPMY01, distance 0.027-0.029, lower than 0.04) and two conspecific strains in Clade IA (CAJQUD01 and JAEUOK01, distance 0.052-0.061, higher than 0.04).

**Figure 2.**
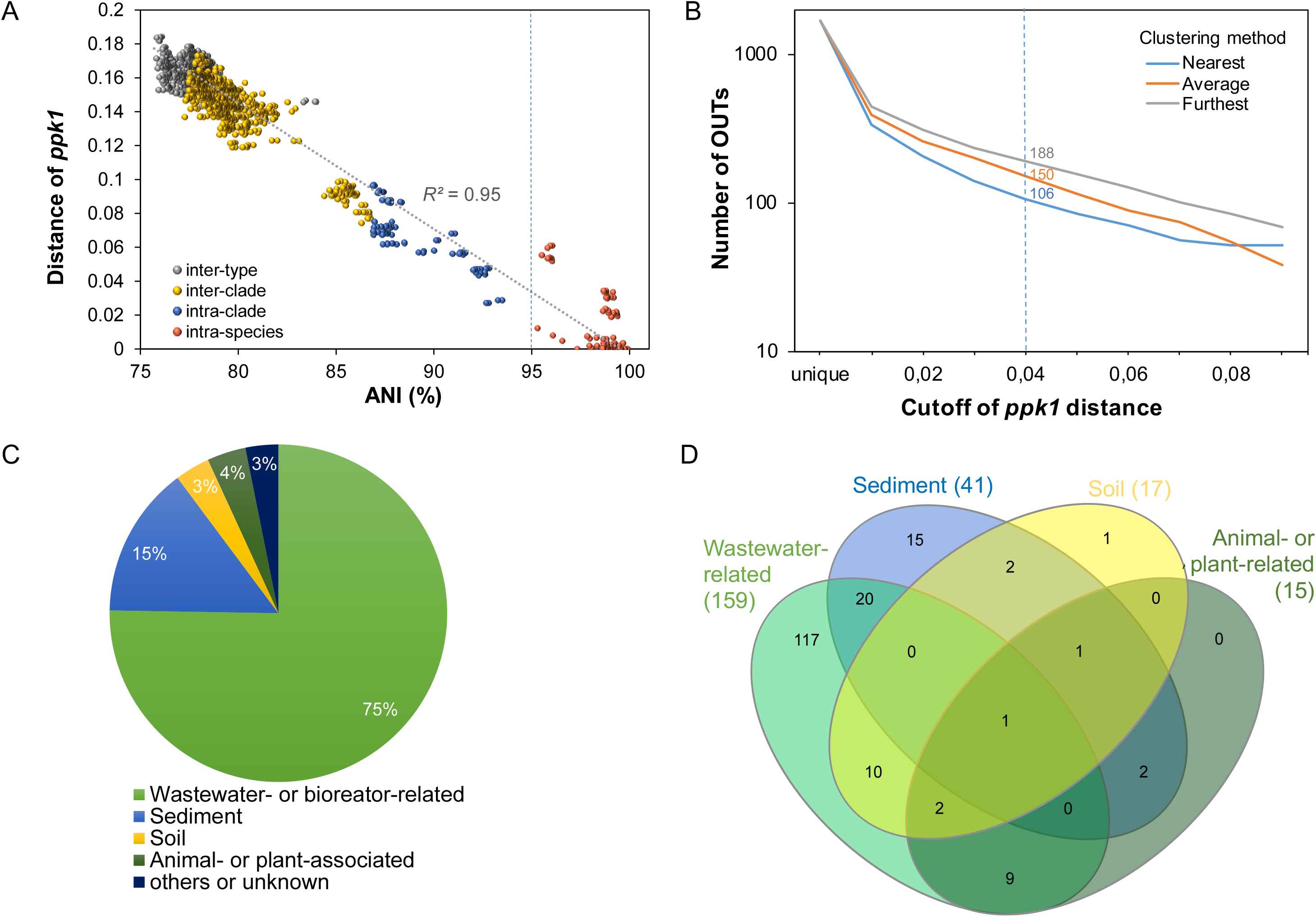
Estimation of the species-level diversity of *Accumulibacter* based on *ppk1* OTU clustering. A) Correlation between genomic ANI and *ppk1* distance. The grey line is the regression of the correlation. ANI of 95% is the species-level cutoff. B) The number of OTUs obtained by *ppk1* clustering with nearest, average, and furthest algorithms under various cutoffs. The 0.04 cutoff is referred to since it was proposed as the species-level threshold. C) Sources of qualified *ppk1* sequences downloaded from Genbank and their percentage. D) Venn diagram showing the distribution of species-level *ppk1* OTUs among four major environmental sources.

A GenBank collection of potential *ppk1* sequences from *Accumulibacter* lineage was cleaned, aligned, trimmed, phylogenetically verified, and dereplicated (See Methods). Finally, 1,665 unique and zero-gap *ppk1* alignments (1,006 sites) were retained for analysis. According to the furthest, average, and nearest OTU clustering, the numbers of OTUs at 0.04 threshold ranged from 106 (the nearest algorithm) to 188 (the furthest algorithm), indicating an unconscious species-level diversity in *Accumulibacter* (Figure 2B). Therefore, the majority of *Accumulibacter* species still has no genome. As displayed in Figure 2C, the sources of the collected *ppk1* sequences from GenBank showed that three-quarters were retrieved from wastewater-related environments, followed by sediment (15%). Twenty-one OTUs (20 from sediments) have not been previously detected in wastewater-related systems (Figure 2D).

### Transcriptional feature among co-occurring *Accumulibacter* strains

Basic information on the two metatranscriptomic data (AN and AE) is listed in Table S5. We performed genome-centered metatranscriptomic analysis on the seven co-occurring *Accumulibacter* strains (Bin142, 208, 136, 28, 228, 87, and 45), each of which has over one million transcriptomic reads (rRNA removed) in both datasets. We determined the relative gene expression level (in RPM, see Materials and Methods) for each gene in each genome. Only key functional gene catalogs related to the EBPR performance and PAO phenotypes, such as denitrification, poly-P-accumulation, and PHA and glycogen metabolism, were examined to decipher the inter-strain transcriptional feature. In most cases, we summed the expression level of all copies of a given functional gene (e.g., Bin 142 has three copies of *phaC*, then *phaC* expression level was calculated by summing up the expression levels of all three copies). However, the multicopy of *Pit* was analyzed independently to examine the inter-copy transcriptional feature.

Figures 3A&B show that during anaerobic and aerobic phases, the inter-strain expression patterns of these PAO phenotype-related genes had minimal correlation with their phylogeny. Although the two Clade IIC strains (Bin142 and Bin28) were clustered first in both phases, the two conspecific Type III strains (Bin228 and Bin87) did not exhibit highly similar expression patterns. The dynamics of gene expression during both phases was examined by calculating the ratio of the relative expression level for a given gene (Figure 3C). Intriguingly, for the co-occurring *Accumulibacter* strains, PHA and glycogen metabolism genes showed less inter-phase dynamics and inter-strain divergences than the genes related to denitrification and poly-P accumulation. The AN-to-AE fold change of gene expression level ranged 0.4–1.9 and 0.2–15.2 for the former two and the latter two, respectively.

**Figure 3.**
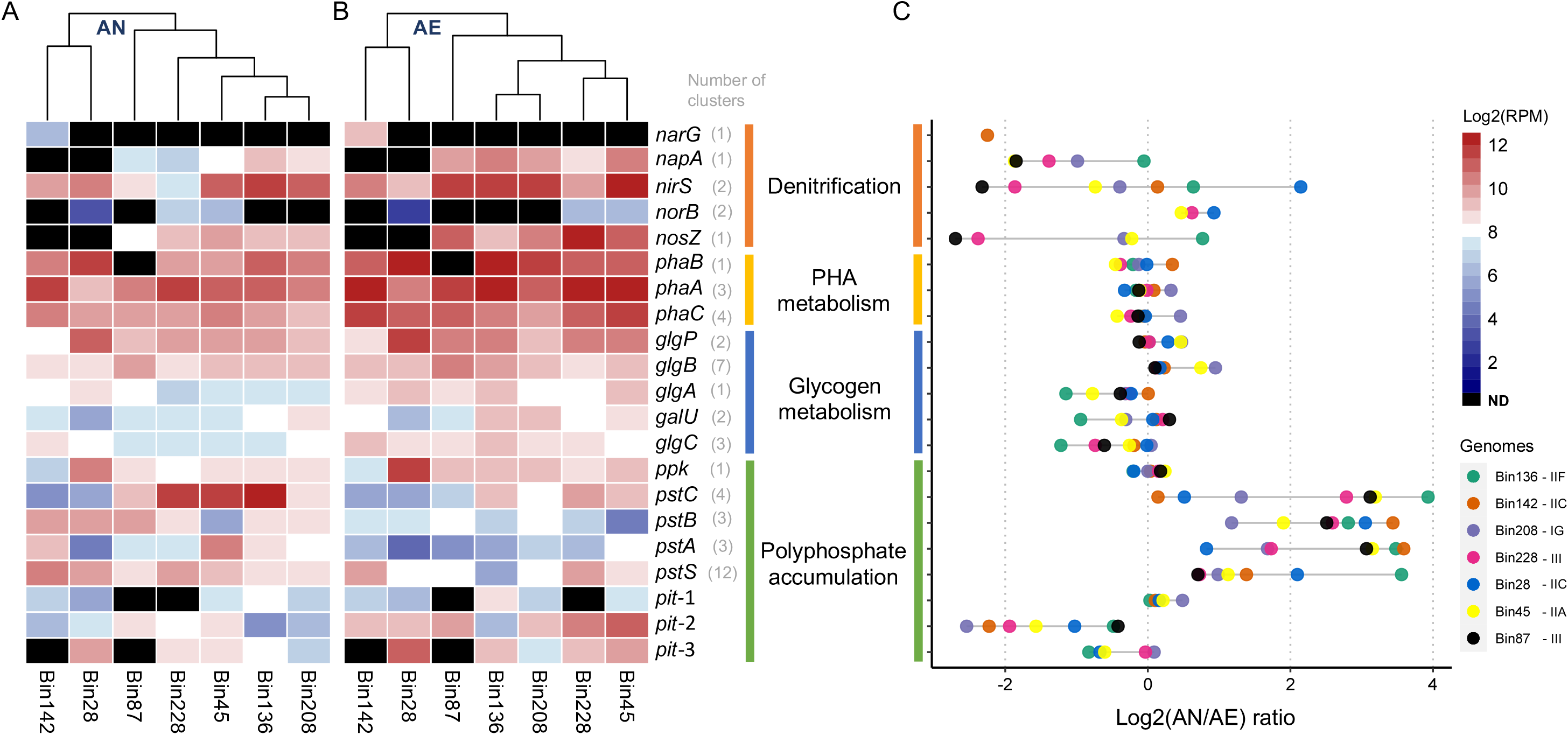
Metatranscriptomic analysis of key functional genes in denitrification, polyphosphate accumulating, PHA, and glycogen metabolism in seven co-occurring *Accumulibacter* strains at AN (A) and AE (B). ND: not detected at the DNA level. The columns were clustered with Euclidean distance. Besides *pit*, the expression level was the sum of all copies for a given gene and was shown in the log2 scale. The copy number of a gene cluster was shown beside the gene names. AN-to-AE ratios of expression levels for all genes and all strains were shown in C.

Among the denitrification genes, *narG*, which is only present in Bin142, showed a higher expression level in AE than in AN, similar to all *napA* genes that perform aerobic nitrate reduction. Although NAR was thought to be suppressed under aerobic conditions, the nitrate level was extremely low (<0.2 mg L^-1^) at the sampling time of AN (Fig S1). Despite the high DO in the bulk water, the anoxic condition may occur in the microenvironment of biomass. Moreover, whether *nirS* and *nosZ* exhibits diverse expression dynamics among the strains remained unclear. Among the poly-P accumulation-related genes, *ppk1* showed the smallest dynamics. The four high-affinity phosphate transporter genes (*pstA, pstB, pstC*, and *pstS*) had a higher expression level in AN than AE, and two of the three *Pit* copies encoding the low-affinity phosphate transporter showed a higher expression level in the AE phase than in AN (Figure 4C). One copy (*Pit*-2) had the highest inter-phase dynamics and inter-strain divergence. The two dominant strains, Bin142 and Bin208, had the highest inter-phase dynamics of this gene copy.

**Figure 4.**
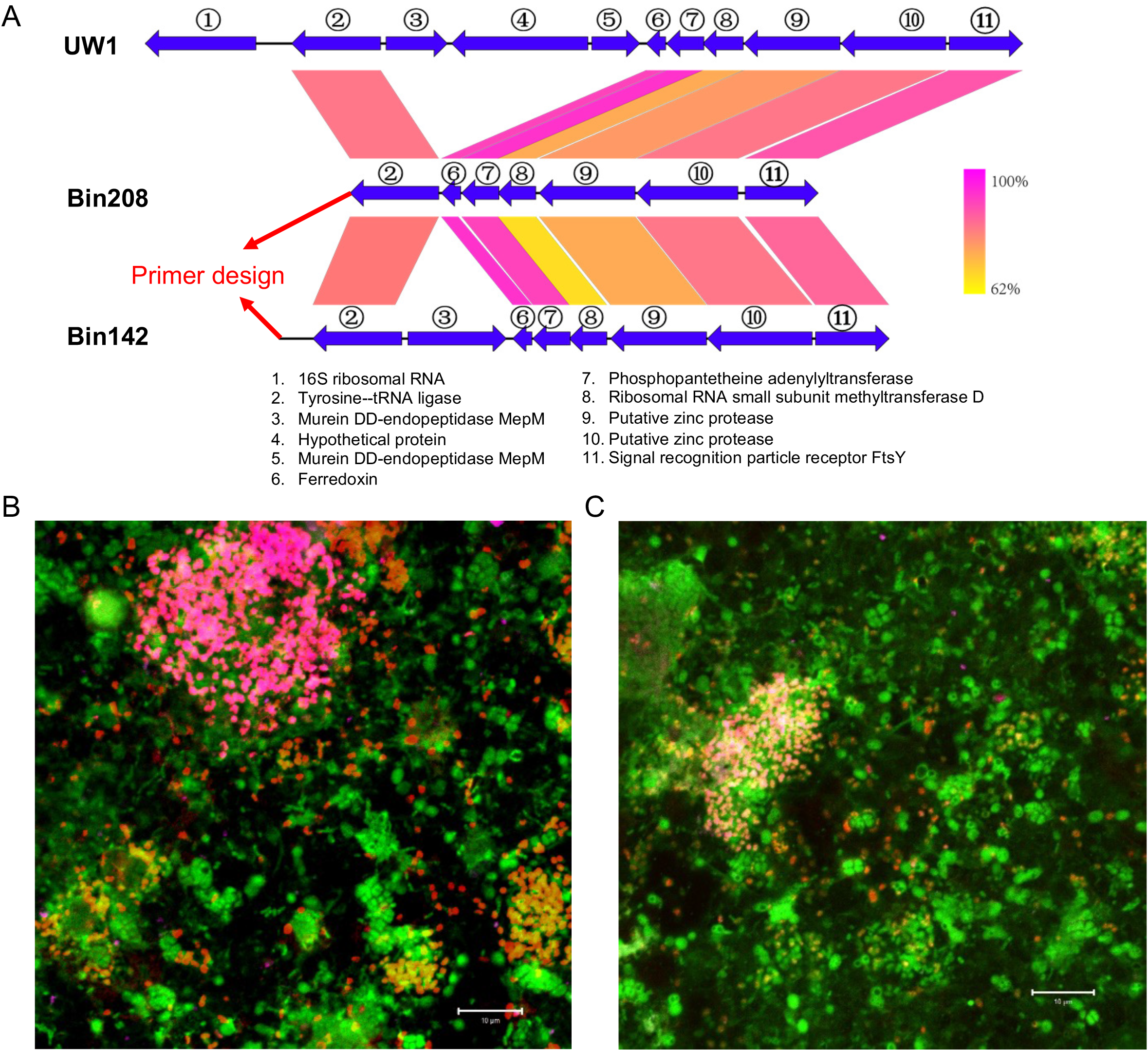
FISH detection of two co-occurring *Accumulibacter* strains (Bin142 and Bin208). Primer design based on the genomic synteny of the upstream region of 16S rRNA genes among the two strains and UW-1 (A). Based on the cloning of 16S rRNA gene, FISH probe was designed for both strains. FISH was performed by using the probes of EUB338-mix (all bacteria, in green), PAO-mix (all *Accumulibacter*, in red), and Bin208 (B) or Bin142 (C), which were labeled in pink. Bar=10 μm.

### Retrieving full-length 16S rRNA and FISH detection of Bin142 and Bin208

Morphologically distinguishing closely related taxa by rRNA-based FISH in activated sludge samples is technically challenging because of the difficulty in obtaining their full-length rRNA sequences in MAGs generated by short-read next-generation sequencing (41). As expected, all 10 *Accumulibacter* bins did not contain the 16S rRNA gene. We applied a strategy based on strain-specific primer design to obtain the full-length sequences of 16S rRNA gene for IG (Bin208) and IIC (Bin142), the top two abundant *Accumulibacter* strains in the bioreactor I sample (accounting for approximately 80% of total *Accumulibacter* according to their genomic coverages, see Fig S3). Given that the FISH samples were collected 19 months after the metagenomic samples, a *ppk1* clone library was constructed to confirm the existence of the two strains (each >30% of total *Accumulibacter*). As Bin142 had no *ppk1* in its genome, the *ppk1* from its conspecific strain was used as reference (Figure 1B).

First, we extracted 10 ORFs upstream of a certain 16S rRNA gene copies in the whole genome of UW-1 and used them as references to identify the homogenous genomic region in Bin208 and Bin142, which were both broken at the upstream region of the putative 16S rRNA gene (Figure 4A). A specific primer putatively close to the downstream 16S rRNA gene was designed for each bin, and PCR products were successfully obtained using the primer and 1492R (Table S6). The full-length 16S rRNA gene was retrieved for both genomes. The two genomes shared 98.7% and 98.4% similarity with UW-1 (accession no: CP001715), and the sequence similarity between them was 98.2%.

We found that the requirement of FISH probe specific for each strain among the domain of bacteria or even only within *Accumulibacter* is impossible to achieve. In addition, we have no rRNA gene sequence information for other co-occurring *Accumulibacter* strains. Thus, we followed a second-best strategy only requiring that the probes can target the corresponding strains considering the two strains are dominant populations (information on FISH probes can be found in Table S6). Simultaneously hybridized with EUB338-mix (targeting nearly all bacteria) and PAO-mix (targeting all *Accumulibacter*) cells respectively targeted Bin208 and Bin142 specific probes, which are both coccoid, similar to other reported *Accumulibacter* strains (17). The size of the former was slightly larger than that of the latter (Figures 4B&C). Both strains formed loose clusters in bioflocs. No morphological divergence was observed for the cells targeted by each FISH probe, suggesting that other co-occurring *Accumulibacter* strains (without knowing their rRNA gene sequence) might not be targeted by the probes or other false-positive targets are morphologically indistinguishable from Bin208 or Bin142.

## Discussion

Although *Accumulibacter* spp. are not the only PAOs in WWTPs (42–44), they are the core taxa of global WWTP microbial consortia (45) and the PAOs with the high per cell poly-P content (46). Metagenomic approach is continuously promoting our understanding of the genomic and functional diversity of *Accumulibacter* (16, 17). In the present study, we contributed novel *Accumulibacter* genomes representing a novel type. By building the correlation between genomic ANI and *ppk1* similarity, we estimated the global species-level diversity of *Accumulibacter* for the first time. Although using a slightly different genomic dataset, Petriglieri et al. (2022) (17) also defined 26 species in *Accumulibacter*, the same as the ANI-deduced species number in the present study. However, our estimation indicated that currently known species with genomes may only represent the tip-of-the-iceberg diversity for this genus. Moreover, our assessment can be an underestimation for at least two apparent reasons. One is that primer mismatch detected in the two Type III genomes can cause the missing of certain lineages in previous studies using the widely adopted *ppk1* primer sets (254F and 1376R) (47). Metagenomic sequencing, or using a revised primer with higher coverage, can solve this problem. The other reason is the sample representativity. Although biomass samples from multiple types of bioreactors and full-scale WWTPs have been extensively examined for the diversity of *Accumulibacter*, the unexamined ones are reasonably high in number. As suggested by previous studies and ours, the sediment may be the natural habitat for the majority of *Accumulibacter* (48, 49). Given that the diversity from the sediment samples is insufficient, further investigations are required via *ppk1* profiling or metagenomic sequencing.

Expanding the phylogenetic and genomic diversity of *Accumulibacter* can also bring new insights into its functional diversity, especially those potentially related to wastewater treatment. It is suggested that all Type III genomes can utilize the organic electron acceptor, DMSO, for energy production. DMSO, together with its chemical relatives, dimethyl sulfide (DMS) and dimethylsulfoniopropionate, is biologically produced in marine and freshwater systems, especially in sediments, mainly through the methylation of hydrogen sulfide and methylmercaptan (50). The ability of Type III *Accumulibacter* strains to encode DMSO reductase also indicates that the sediment is the natural habitat of certain *Accumulibacter* species. Furthermore, this novel functional feature in Type III may be meaningful for EBPR in certain wastewater systems. As a versatile solvent, DMSO is widely applied in various industries (51). Although its removal can be a biotechnological bottleneck (52), it only degrades after being reduced to DMS (53). Type III *Accumulibacter* may be the candidate treating organisms for treating wastewater containing high concentrations of DMSO and phosphate.

Although the central metabolic model for the PAO phenotype of *Accumulibacter* was established mainly based on the UW-1 strain, the genetic feature that determined this phenotype remains unclear. What are the genes and regulatory characteristics determining *Accumulibacter* spp. becoming PAOs in EBPR systems? With the continuous accumulation of genomic information for *Accumulibacter*, other PAOs, such as the two newly verified ones affiliated with *Dechloromonas* and *Ca*. Phosphoribacter (formerly *Tetrasphaera*) (42, 43) and their phylogenetic relatives without PAO phenotypes, this question can be answered to some extent. Here, we discussed *Pit*, a gene feature unique to PAOs compared with their EBPR competitors, GAOs (54). Our comparative genomics indicated that outgroups of *Accumulibacter* also contain this gene and the divergence between *Accumulibacter* and outgroups is copy number (Figure 1B). Intriguingly, *Pit* also exhibited a higher copy number in the genomes of PAOs affiliated with *Dechloromonas* and *Ca*. Phosphoribacter than in those of the corresponding outgroups (four copies in a representative *Dechloromonas* PAO genome, not present in the study (42), determined by us; 2-5 and 1-2 copies in *Ca*. Phosphoribacter and its outgroups (43)). The multi-copies of *Pit* seem to be a link to the divergent regulation feature. Metatranscriptomic results indicated that only one copy of *Pit* exhibited a predicted P transportation model (i.e., P-influx in the aerobic phase); the other copies had no consistent regulatory feature and were typically expressed at a low level (Figure 3). An early metaproteomic investigation of the EBPR system found that different *pit* proteins of *Accumulibacter* have divergent dynamics between the two phases (55). In the present and that studies, the high-affinity inorganic phosphate transporter (PST) consistently exhibited high expression levels in the anaerobic phase. This finding raised the possibility that P efflux in the anaerobic phase may be facilitated by the PST system.

Among the co-occurring *Accumulibacter* strains, the inter-phase expression dynamics of key genes involved in denitrification and phosphate accumulation showed higher inter-strain divergence than those involved in carbon metabolism (Figure 3). Previous studies also reported the high divergence of the N and P metabolism gene expression pattern among co-occurring *Accumulibacter* (6, 18). These consistent results suggested that different *Accumulibacter* strains may have different niche adaptative features and capabilities in denitrification and phosphate accumulation, no matter if their gene composition is the same. For example, the sole *Accumulibacter* strain with a complete denitrification pathway, QPGA, only highly expresses denitrifying genes under the microaerobic condition potentially via the regulation of FNR regulon (56). Moreover, since seven strains representing multiple clades and types can be convincingly solved by genome-centered metatranscriptomics in our datasets, the gene expression pattern can be linked to their phylogeny. Unexpectedly, the results indicated the decoupling manner between phylogeny and expression pattern of those key genes involved in EPBR functions. Therefore, type-, clade-, or even species-level profiling may not be a confident basis for predicting gene expression patterns and function. This result reminded us of an additional example that *Accumulibacter* strains within the same clade can exert both PAO and GAO phenotypes under the same bioreactor setting (57).

Different *Accumulibacter* species seem to have similar coccoid morphology, despite being different in cell size (17). FISH is perhaps the only option to determine the cell morphology of uncultured microorganisms in a complex community. Petriglieri et al. (2022) (17) performed metagenomic sequencing using long-read technology and retained rRNA genes in the assembled genomes with a high chance. Our pipeline gives another feasible way for cloning rRNA sequences for metagenomic bins having phylogenetic relatives with complete genomes. Cell morphology and size contain key information for the estimation of the population-level contribution of poly-P-accumulating capability. Combining the population-level FISH with the poly-P measurement approach, such as Raman microspectroscopy, can quantitatively assign the contribution of each population (46).

## Conclusions

In this study, the phylogenetic and genomic diversity of *Accumulibacter* was expanded, and one novel type (Type III) was proposed for the first time. The findings suggested that the species-level diversity in this key genus of EPBR is so diverse, and only the tip of the iceberg has been discovered so far. Continuously expanding our understanding of its genomic and functional diversity may be critical for various biotechnological applications, such as the potential DMSO-reducing capability of Type III genomes. With accumulating data on related genomes and performance, the genetic foundation for PAO phenotype in EPBR systems can be further explored and used in predicting unknown but promising PAO candidates. On the basis of our results, the functional feature of each *Accumulibacter* strain may not be necessarily related to the phylogeny, such as type, clade, and even species. Deciphering the function and capability of certain *Accumulibacter* should be specific to the environment (i.e., settings in a bioreactor or full-scale WWTP) and population.

## Data availability

The Illumina short-read sequence datasets have been deposited under the NCBI BioProjects PRJNA524249 (metagenomic data with the accession numbers SRR8926552 and SRR8926553) and PRJNA892277 (metatranscriptomic data and genome bins).

## Acknowledgement

This study was funded by the National Natural Science Foundation of China (No. 31500100) and Innovation Group Project of Southern Marine Science and Engineering Guangdong Laboratory (Zhuhai) (No. 311021006).

## Figure captions

Fig S1. Typical operational parameters during one EBPR cycle. The concentration of COD, PO_4_-P, and NH_4_-N are measured as 635.1, 27.8, and 24.4 mg/L in the raw artificial wastewater. Aeration was started at 120 min. Sampling times are at 5, 15, 35, 55, 85, 110, 130, 145, 175, 205, 235, 265, 325 and 350 minutes. The two metatransciprtomic samples were collected at 15 and 130 minutes, respectively.

Fig S2. Pairwise ANI of 10 *Accumulibacter* bins extracted from metagenomes from this study. ANI values were shown in the heatmap. Left side was the phylogenetic tree based on bac120 genes.

Fig S3. Genome coverage (per GB) of 10 *Accmulibacter* bins in the two metagenomes.

Fig S4. Primers used to amplify the *ppk1* fragment of *Accumulibacter* showed mismatches in Bin87 and Bin228 which belong to Type III newly proposed in this study.

Fig S5. Neighbor-joining phylogenetic tree based on globally collected *ppk1* representatives. The *ppk1* sequences of the 0.09-OTU representatives and those extracted from genomes were involved. Bootstrap values were shown beside the nodes of types and clades. The *ppk1* sequences from two *Propionivibrio* genomes were used as outgroups.

Table S1. Information on metagenomic bins of *Accumulibacter*, Competibacteraceae, and other taxa.

Table S2. Information on downloaded reference genomes

Table S3. The lost genes in Type III *Accumulibacter* genomes

Table S4. Distribution of key genes in PAO phenotype

Table S5. Basic information on the two metatranscriptomic datasets

Table S6. Primers and probes used in the study

